# Multi-ethnic polygenic risk scores improve risk prediction in diverse populations

**DOI:** 10.1101/051458

**Authors:** Carla Márquez-Luna, Po-Ru Loh, South Asian Type 2 Diabetes (SAT2D) Consortium, The SIGMA Type 2 Diabetes Consortium, Alkes L. Price

## Abstract

Methods for genetic risk prediction have been widely investigated in recent years. However, most available training data involves European samples, and it is currently unclear how to accurately predict disease risk in other populations. Previous studies have used either training data from European samples in large sample size or training data from the target population in small sample size, but not both. Here, we introduce a multi-ethnic polygenic risk score that combines training data from European samples and training data from the target population. We applied this approach to predict type 2 diabetes (T2D) in a Latino cohort using both publicly available European summary statistics in large sample size and Latino training data in small sample size. We attained a >70% relative improvement in prediction accuracy (from *R*^2^=0.027 to *R*^2^=0.047) compared to methods that use only one source of training data, consistent with large relative improvements in simulations. We observed a systematically lower load of T2D risk alleles in Latino individuals with more European ancestry, which could be explained by polygenic selection in ancestral European and/or Native American populations. Application of our approach to predict T2D in a South Asian UK Biobank cohort attained a >70% relative improvement in prediction accuracy, and application to predict height in an African UK Biobank cohort attained a 30% relative improvement. Our work reduces the gap in polygenic risk prediction accuracy between European and non-European target populations.

**Author Summary:** The use of genetic information to predict disease risk is of great interest because of its potential clinical application. Prediction is performed via the construction of polygenic risk scores, which separate individuals into different risk categories. Polygenic risk scores can also be applied to improve our understanding of the genetic architecture of complex diseases. The ideal training data set would be a large cohort from the same population as the target sample, but this is generally unavailable for non-European populations. Thus, we propose a summary statistics based polygenic risk score that leverages both a large European training sample and a training sample from the same population as the target population. This approach produces a substantial relative improvement in prediction accuracy compared to methods that use a single training population when applied to predict type 2 diabetes in a Latino cohort, consistent with simulation results. We observed similar relative improvements in applications to predict type 2 diabetes in a South Asian cohort and height in an African cohort.

## Introduction

Genetic risk prediction is an important and widely investigated topic because of its potential clinical application as well as its application to better understand the genetic architecture of complex traits [1]. Many polygenic risk prediction methods have been developed and applied to complex traits. These include polygenic risk scores (PRS) [2–9], which use summary association statistics as training data, and Best Linear Unbiased Predictor (BLUP) methods and their extensions [10–17], which require individual-level genotype and phenotype data.

However, all of these methods are inadequate for polygenic risk prediction in non-European populations, because they consider training data from only a single population. Existing training data sets have much larger sample sizes in European populations, but the use of European training data for polygenic risk prediction in non-European populations reduces prediction accuracy, due to different patterns of linkage disequilibrium (LD) in different populations [2,8,18,19]. For example, ref. [8] reported a relative decrease of 53-89% in schizophrenia risk prediction accuracy in Japanese and African-American populations compared to Europeans when applying PRS methods using European training data. An alternative is to use training data from the same population as the target population, but this would generally imply a much lower sample size, reducing prediction accuracy.

To tackle this problem, we developed an approach that combines PRS based on European training data with PRS based on training data from the target population. The method takes advantage of both the accuracy that can be achieved with large training samples [4,5] and the accuracy that can be achieved with training data containing the same LD patterns as the target population. In simulations and application to predict type 2 diabetes (T2D) in Latino target samples in the SIGMA T2D data set [20], we attained a >70% relative improvement in prediction accuracy (from *R*^2^=0.027 to *R*^2^=0.047) compared to methods that use only one source of training data. We also obtained a >70% relative improvement in an analysis to predict T2D in a South Asian UK Biobank cohort, and a 30% relative improvement in an analysis to predict height in an African UK Biobank cohort.

## Materials and Methods

### Polygenic risk score using a single training population

Polygenic risk scores are constructed using SNP effect sizes estimated from genome-wide association studies, which perform marginal regression of the phenotype of interest on each SNP in turn. Explicitly, for continuous traits, we estimate effect sizes 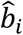 (where *i* = 1,…,*M* indexes genetic markers) using the model *y* = *b*_0_ + *b_i_g_i_* + *b_PC_PC* + *ε*, where *g_i_* denotes genotypes at marker *i*, *PC* denotes one or more principal components used to adjust for ancestry, and ε denotes environmental noise. For binary traits, we use the analogous logistic model logit[P(*y*=1)] = *b*_0_ + *b_i_g_i_* + *b_PC_PC* + *ε*.

Given a vector of estimated effect sizes 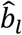 from a genome-wide association study performed on a set of training samples, the polygenic risk score [2] (PRS) for a target individual with genotypes *g_i_* is defined as 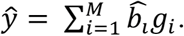 In practice, rather than computing the PRS using estimated effect sizes for all available genetic markers, the PRS is computed on a subset of genetic markers obtained via informed LD-pruning [3] (also known as LD-clumping) followed by P-value thresholding [2]. Specifically, this “pruning + thresholding” strategy has two parameters, *R_LD_*^2^ and *P_T_*, and proceeds as follows. First, we prune the SNPs based on a pairwise threshold *R_LD_*^2^, removing the less significant SNP in each pair (using PLINK; see Web Resources). Second, we restrict to SNPs with an association P-value below the significance threshold *P_T_*.

The parameters *R_LD_*^2^ and *P_T_* are commonly tuned using on validation data to optimize prediction accuracy [2,3]. While in theory this procedure is susceptible to overfitting, in practice, validation sample sizes are typically large, and *R_LD_*^2^ and *P_T_* are selected from a small discrete set of parameter choices, so overfitting is considered to have a negligible effect. Accordingly, in this work, we consider *R_LD_*^2^ ∈ {0.1, 0.2, 0.5, 0.8} and *P_T_* ∈ {1.0, 0.8, 0.5, 0.4, 0.3, 0.2, 0.1, 0.08, 0.05, 0.02, 0.01, 10^−3^, 10^−4^, 10^−5^, 10^−6^, 10^−7^, 10^−8^}, and we always report results corresponding to the best choices of these parameters. In all of our primary analyses involving two training populations (see below), values of *R_LD_*^2^ and *P_T_* were optimized based only on PRS in a single training population, to ensure that PRS using two training populations did not gain any relative advantage from the optimization of these parameters.

In this work, we specifically consider PRS built using European (EUR), Latino (LAT), South Asian (SAS), or African (AFR) training samples. We use the notation *PRS_EUR_* to denote PRS built using European samples, and analogously for the other populations.

### Polygenic risk score using two training populations

Given a pair of polygenic risk scores computed as above using two distinct training populations, we define the multi-ethnic PRS with mixing weights *α_1_* and *α_2_* as the linear combination of the two PRS with these weights: e.g., for EUR and LAT, we define *PRS_EUR+LAT_ = α_1_PRS_EUR_ + α_2_PRS_LAT_*. We employ two different approaches to avoid overfitting. In our primary analyses, we estimate mixing weights *α_1_* and *α_2_* using validation data and compute adjusted *R*^2^ to account for the additional degree of freedom. In our secondary analyses, we estimate mixing weights *α_1_* and *α_2_* using cross-validation (see Assessment of methods below).

### Polygenic risk score using one or two training populations and genetic ancestry

We further define polygenic risk scores that include an ancestry predictor, namely, the top principal component in a given data set. (We considered only the top PC in each data set that we analyzed, because lower PCs had a squared correlation with phenotype lower than 0.005 in each case.) We define a polygenic risk score LAT+ANC with mixing weights *α_1_* and *α_2_* as *PRS_LAT+ANC_ = α_1_PRS_LAT_ + α_2_ PC*, and we define a polygenic risk score EUR+LAT+ANC with mixing weights *α_1_*, *α_2_* and *α_3_* as *PRS_EUR+LAT+ANC_ = α_1_PRS_EUR_ + α_2_PRS_LAT_ + α_3_PC*. As above, we employ two different approaches to avoid overfitting: in our primary analyses, we estimate mixing weights using validation data and compute adjusted *R*^*2*^; in our secondary analyses, we estimate mixing weights using cross-validation.

### Assessment of methods

We assessed the accuracy of polygenic risk scores in validation samples (independent from samples used to estimate effect sizes). We used adjusted *R*^2^ as the accuracy metric for continuous traits and liability-scale adjusted *R*^2^ (ref. [21]) for binary traits. Adjusted *R*^2^ is defined as 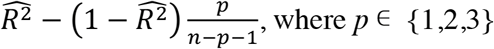 is the number of PRS or ANC components in the mixture, *n* is the number of validation samples, and 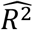 is the raw (unadjusted) *R*^2^. The adjusted *R*^2^ metric roughly corrects for increased model complexity in multi-component PRS, so in our primary analyses, we report accuracy as adjusted *R*^2^ using best-fit mixing weights 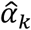 estimated using the validation data.

To verify that this metric provides robust model comparisons, we also performed auxiliary analyses in which we used 10-fold cross-validation: specifically, for each left-out fold in turn, we estimated mixing weights using the other 9 folds and evaluated adjusted *R*^2^ for PRS computed using these weights on the left-out fold. We then computed average adjusted *R*^2^ across the 10 folds. (When analyzing data from an unbalanced case-control study with #cases << #controls, we used stratified 10-fold cross-validation, selecting the folds such that each fold had the same case-control ratio; this applies only to the South Asian UK Biobank T2D analysis.)

Finally, for analyses in which we needed to use samples from the same cohort for both building PRS (i.e., estimating effect sizes 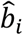) and validation, we also used cross-validation. In our primary analyses, we employed 10-fold cross-validation, using 90% of the cohort to estimate 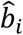 and the remaining 10% of the cohort to validate predictions (using the adjusted *R*^2^ metric with best-fit mixture weights 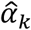). In our secondary analyses, we employed 10×9-fold cross-validation, in which 90% of the cohort was used to estimate both 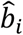 and 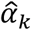 and the remaining 10% of the cohort was used to validate predictions. To estimate 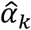, we iteratively split the 90% set of training samples into an 80% training-training set and a 10% training-test set; we estimated 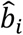 in the 80% training-training set and computed a PRS for the 10% training-test set for each of the 9 training-test folds, and we then performed a single regression of phenotype against each PRS across the entire 90% set of training samples to estimate 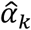. Finally, we reestimated 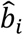 for the final test prediction using the entire 90% set of training samples.

### Simulations

We simulated quantitative phenotypes using real genotypes from European (WTCCC2) and Latino (SIGMA) data sets (see below). We fixed the proportion of causal markers at 1% and fixed SNP-heritability *h_g_*^2^ at 0.5, and sampled normalized effect sizes *β_i_* from a normal distribution with variance equal to *h_g_*^2^ divided by the number of causal markers. We calculated per-allele effect sizes *b_i_* as 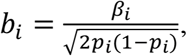 where *p_i_* is the minor allele frequency of SNP *i* in the European data set. We simulated phenotypes as 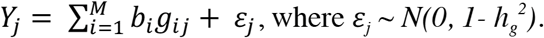

In our primary simulations, we discarded the causal SNPs and used only the non-causal SNPs as input to the prediction methods (i.e. we simulated untyped causal SNPs, which we believe to be realistic). As an alternative, we also considered simulations in which we included the causal SNPs as input to the prediction methods (i.e., a scenario in which causal SNPs are typed).

We also performed simulations in which Latino phenotypes were explicitly correlated to ancestry (population stratification). In these simulations, we added a constant multiple of PC1 (representing European vs. Native American ancestry, with positive values representing higher European ancestry) to the Latino phenotypes such that the correlation between phenotype and PC1 was equal to -0.11, which is the correlation between the T2D phenotype and PC1 in the SIGMA data set.

We performed simulations under 4 different scenarios: (i) using all chromosomes, (ii) using chromosomes 1-4, (iii) using chromosomes 1-2, and (iv) using chromosome 1 only. The motivation for performing simulations with a subset of chromosomes was to increase *N*/*M*, extrapolating to performance at larger sample sizes, as in previous work [8].

### Simulation data sets: WTCCC2 and SIGMA

Our simulations used real genotypes from the WTCCC2 and SIGMA data sets. The WTCCC2 data set consists of 15,622 unrelated European samples from a multiple sclerosis study genotyped at 360,557 SNPs after QC [22,23] (see Web Resources). The SIGMA data set consists of 8,214 unrelated Latino samples genotyped at 2,440,134 SNPs after QC [20] (see Web Resources). We restricted our simulations to 232,629 SNPs present in both data sets (with matched reference and variant alleles) after removing A/T and C/G SNPs to eliminate potential strand ambiguity.

### Latino type 2 diabetes data sets: DIAGRAM, SIGMA and UK Biobank

Our analyses of type 2 diabetes in Latinos used summary association statistics from the DIAGRAM data set and genotypes and phenotypes from the SIGMA data set. The DIAGRAM data set consists of 12,171 cases and 56,862 controls of European ancestry for which summary association statistics at 2,473,441 imputed SNPs are publicly available (see Web Resources) [24]. As noted above, the SIGMA data set consists of 8,214 unrelated Latino samples (3,848 type 2 diabetes cases and 4,366 controls) genotyped at 2,440,134 SNPs after QC. SIGMA association statistics were computed with adjustment for 2 PCs, as in ref. [20]. We restricted our analyses of type 2 diabetes to 776,374 SNPs present in both data sets (with matched reference and variant alleles) after removing A/T and C/G SNPs to eliminate potential strand ambiguity. For the SIGMA data set, we used the top 2 PCs as computed in ref. [20]. We also performed an analysis of type 2 diabetes using imputed genotypes from the SIGMA T2D data set [20], restricting to 2,062,617 SNPs present in both data sets (with matched reference and variant alleles) after removing A/T and C/G SNPs to eliminate potential strand ambiguity.

We performed a secondary analysis using 113,851 British samples from UK Biobank [25] (see Web Resources) as European training data (5,198 type 2 diabetes cases and 108,653 controls). UK Biobank association statistics were computed with adjustment for 10 PCs [25], estimated using FastPCA [26] (see Web Resources). We computed summary statistics for 608,878 genotyped SNPs from UK Biobank after removing A/T and C/G SNPs to eliminate potential strand ambiguity. We analyzed 187,142 SNPs present in the SIGMA and UK Biobank data sets. We defined type 2 diabetes cases in UK Biobank as “any diabetes” with “age of diagnosis > 30”. We note that the p-values at two top type 1 diabetes (T1D) loci (rs2476601, rs9268645) were only nominally significant (p~0.05) for this T2D phenotype, indicating low contamination with T1D cases.

### South Asian type 2 diabetes data sets: DIAGRAM, SAT2D and UK Biobank

Our analysis of type 2 diabetes in South Asians used European summary association statistics from the DIAGRAM data set (described above), South Asian summary statistics data from the South Asian Type 2 Diabetes (SAT2D) Consortium [27], and South Asian genotypes and phenotypes from UK Biobank (see Web Resources) as test data. The SAT2D data set consists of 5,561 South Asian type 2 diabetes cases and 14,458 South Asian controls for which we summary statistics for 2,646,472 imputed SNPs were available. The UK Biobank test data consists of 1,756 unrelated samples of South Asian ancestry (272 type 2 diabetes cases and 1,484 controls), genotyped at 608,878 SNPs after QC, with the following self-reported ethnicity distribution: 52 Bangladeshi, 1,301 Indian and 403 Pakistani. We analyzed 208,400 SNPs present in the DIAGRAM, SAT2D and UK Biobank data sets after removing A/T and C/G SNPs to eliminate potential strand ambiguity.

### African height data sets: UK Biobank and N’Diaye et al

Our analyses of height in Africans used European summary association statistics from UK Biobank (see Web Resources), African summary statistics from N’Diaye et al. [28] and African genotypes and phenotypes from UK Biobank. European summary statistics from UK Biobank were computed using 113,660 British samples for which height phenotypes were available with adjustment for 10 PCs [25], estimated using FastPCA [26] (see Web Resources). The N’Diaye et al. [28] data set consists of 20,427 samples of African ancestry with summary association statistics at 3,254,125 imputed SNPs. The UK Biobank data set consists of 1,745 unrelated samples of African ancestry, genotyped at 608,878 SNPs after QC, with the following self-reported ethnicity distribution: 743 African, 1,002 Caribbean. We restricted our analysis to 232,182 SNPs present in the UK Biobank and N’Diaye et al. data sets after removing A/T and C/G SNPs to eliminate potential strand ambiguity.

## Results

### Simulations

We performed simulations using real genotypes and simulated phenotypes. We simulated continuous phenotypes under a non-infinitesimal model with 1% of markers chosen to be causal with the same effect size in all samples and SNP-heritability *h_g_*^2^ = 0.5 (see Methods); we report the average adjusted *R*^2^ and standard errors over 100 simulations. We used WTCCC2 [22,23] data (15,622 samples after QC; see Methods) as the European training data, and the SIGMA data [20] (8,214 samples) as the Latino training and validation data (with 10-fold cross-validation). We simulated phenotypes using the 232,629 SNPs present in both data sets and built predictions from these SNPs excluding the causal SNPs, modeling the causal SNPs as untyped (see Methods).

Prediction accuracies (adjusted *R*^2^) and optimal weights for the 5 main methods (EUR, LAT, LAT+ANC, EUR+LAT, EUR+LAT+ANC) are reported in Table 1. In each case, the best prediction accuracy was attained using LD-pruning threshold *R*_*LD*_^2^=0.8 (results using different LD-pruning thresholds are reported in S1 Table); the median value of the optimal P-value threshold *P_T_* was equal to 0.01 for EUR and 0.05 for LAT. On average, the EUR method performed only 23% better than the LAT method, despite having twice as much training data. This reflects a tradeoff between the larger training sample size for EUR and the target-matched LD patterns for LAT. EUR+LAT attained 64%-101% relative improvements vs. EUR and LAT respectively (and used a slightly larger weight for EUR than for LAT), highlighting the advantages of incorporating multiple sources of training data. When including an ancestry predictor, EUR+LAT+ANC attained a 10% relative improvement vs. EUR+LAT (≥80% relative improvement vs. EUR or LAT), reflecting small genetic effects of ancestry on phenotype that can arise from random genetic drift between populations at causal markers (which is better-captured by ancestry components than by SNPs used in a PRS). Predictions using Latino effect sizes that were not adjusted for genetic ancestry (LAT_unadj_, EUR+LAT_unadj_, EUR+LAT_unadj_+ANC, as compared to LAT, EUR+LAT, EUR+LAT+ANC) were much less accurate (S2 Table), as in previous work [29]; this is consistent with the fact that LAT_unadj_ predictions were dominated by genetic ancestry (adjusted *R*^2^ = 0.37; S3 Table). We also observed a modest correlation (adjusted *R*^2^ = 0.025) between the EUR prediction and genetic ancestry (S3 Table), again reflecting small genetic effects of ancestry on phenotype that can arise from random genetic drift between populations at causal markers. The relative performance of the different prediction methods was similar in simulations in which phenotypes explicitly contained an ancestry term, representing environmentally-driven stratification (S4 Table).

**Table 1.** Accuracy of 5 prediction methods in simulations. We report average adjusted *R*^2^ over 100 simulations for each of the 5 main prediction methods. We also report normalized weights, defined as the mixing weight 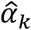 (see Methods) multiplied by the standard deviation of the PRS.

| Model | Average weight (s.e.)<br>associated to each predictor | | Average adj.<br>$R^2$ (s.e.) |
| --- | --- | --- | --- |
|  | European<br>PRS | Latino PRS |  |
| EUR | 0.19449<br>(0.004) |  | 0.03927<br>(0.002) |
| LAT |  | 0.17780<br>(0.003) | 0.03200<br>(0.001) |
| LAT+ANC |  | 0.17613<br>(0.002) | 0.04115<br>(0.002) |
| EUR+LAT | 0.17847<br>(0.004) | 0.15784<br>(0.003) | 0.06441<br>(0.002) |
| EUR+LAT+ANC | 0.19098<br>(0.004) | 0.15578<br>(0.002) | 0.07053<br>(0.002) |

We extrapolated the results in Table 1 to larger sample sizes by limiting the simulations to subsets of chromosomes, as in previous work [8] (Fig 1 and S5 Table). EUR+LAT+ANC was the best performing method in each of these experiments. We also performed simulations using predictions constructed using all SNPs including the causal SNPs (S1 Fig and S6 Table). In these experiments, EUR+LAT+ANC was once again the best performing method, and EUR performed much better than LAT, consistent with the larger training sample size for EUR and the fact that differential tagging of causal SNPs is of reduced importance when causal SNPs are typed.

**Fig 1.**
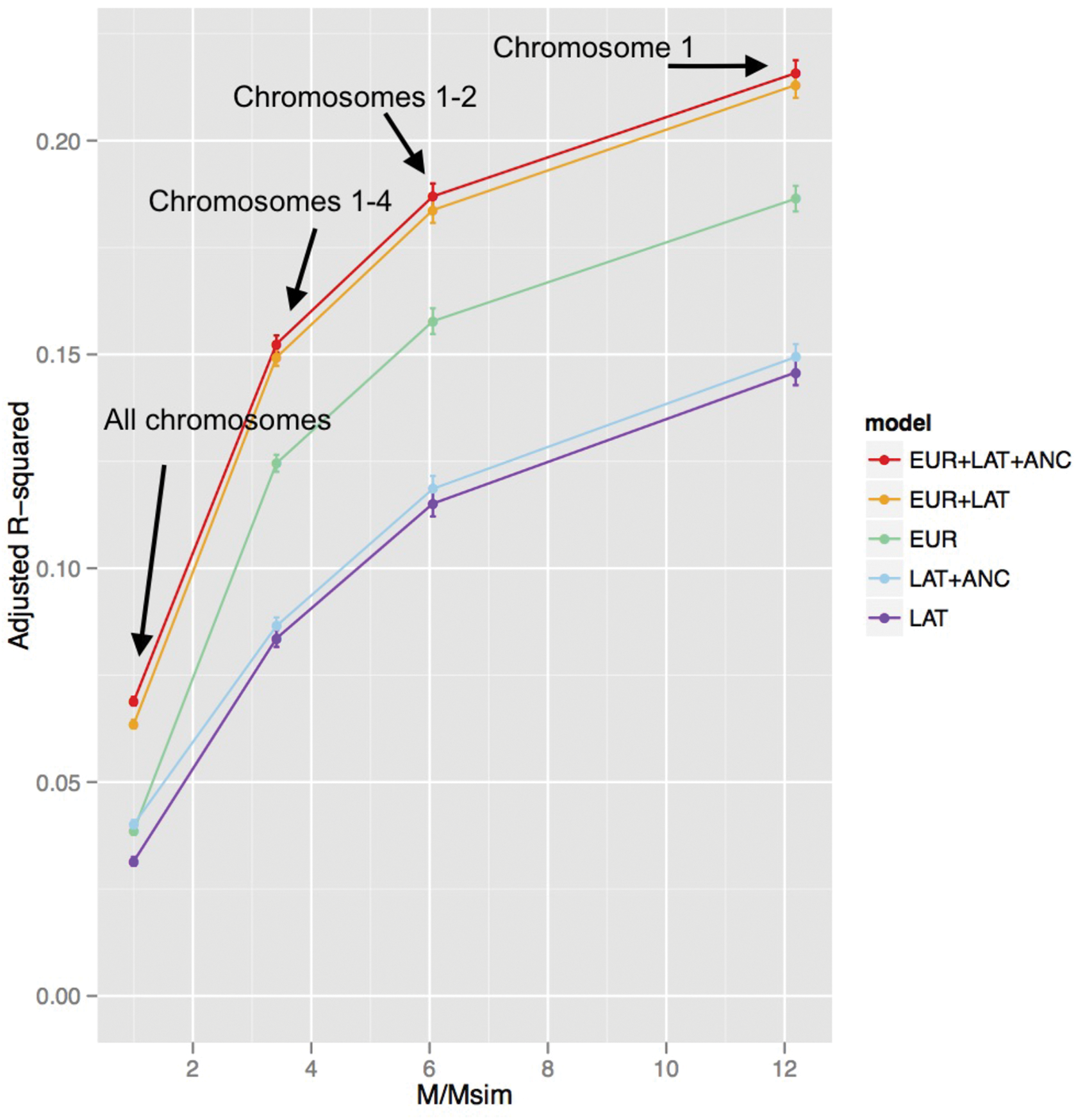
Accuracy of 5 prediction methods in simulations using subsets of chromosomes. We report prediction accuracies for each of the 5 main prediction methods as a function of M/Msim, where M=232,629 is the total number of SNPs and Msim is the actual number of SNPS used in each simulation: 232,629 (all chromosomes), 68,188 (chromosomes 1-4), 38,412 (chromosomes 1-2), and 19,087 (chromosome 1). Numerical results are provided in S5 Table.

### Analyses of type 2 diabetes in Latinos

We applied the same methods to predict T2D in Latino target samples from the SIGMA T2D data set. We used publicly available European summary statistics from DIAGRAM[24] (12,171 cases and 56,862 controls; effective sample size = 4/(1/*N*_case_ + 1/*N*_control_) = 40,101) as European training data and SIGMA T2D genotypes and phenotypes [20] (3,848 cases and 4,366 controls; effective sample size = 8,181) as Latino training and validation data, employing 10-fold cross-validation.

Prediction accuracies (adjusted *R*^2^ on the liability scale [21], assuming 8% prevalence [3]) and optimal weights for the 5 main methods (EUR, LAT, LAT+ANC, EUR+LAT, EUR+LAT+ANC) are reported in Table 2 (other prediction metrics are reported in S7 Table). In each case, the best prediction accuracy was obtained using LD-pruning threshold *R*_*LD*_^2^=0.8 (results using different LD-pruning thresholds are reported in S8 Table); the value of the optimal P-value threshold *P*_*T*_ was equal to 0.05 for EUR and 0.2 for LAT. EUR performed only 33% better than LAT despite the much larger training sample size, again reflecting a tradeoff between sample size and target-matched LD patterns. EUR+LAT attained 75%-133% relative improvements vs. EUR and LAT respectively (and used a slightly larger weight for EUR than for LAT), again highlighting the advantages of incorporating multiple sources of training data. Although adding an ancestry predictor to LAT produced a substantial improvement (LAT+ANC vs. LAT), adding an ancestry predictor to EUR+LAT produced an insignificant change in accuracy for EUR+LAT+ANC compared to EUR+LAT; this can be explained by the large negative correlation between the European PRS (EUR) and the proportion of European ancestry within Latino samples (*R* = −0.75; S9 Table), such that any predictor that includes EUR already includes effects of genetic ancestry. This correlation is far larger than analogous correlations due to random genetic drift in our simulations (S3 Table), suggesting that this systematically lower load of T2D risk alleles in Latino individuals with more European ancestry could be due to polygenic selection [30,31] in ancestral European and/or Native American populations; previous studies using top GWAS-associated SNPs have also reported continental differences in genetic risk for T2D [32,33]. We observed a similar correlation (*R*= −0.77) when using British UK Biobank type 2 diabetes samples as European training data (see Methods), confirming that this negative correlation is not caused by population stratification in DIAGRAM. As in our simulations, predictions using Latino effect sizes that were not adjusted for genetic ancestry (LAT_unadj_, EUR+LAT_unadj_, EUR+LAT_unadj_+ANC, as compared to LAT, EUR+LAT, EUR+LAT+ANC) were much less accurate (S10 Table), consistent with the fact that these predictions were dominated by genetic ancestry (S9 Table). We also computed predictions for each method using imputed SNPs from the SIGMA T2D data set; this did not improve prediction accuracy, but predicting using two training populations still achieved the highest accuracy (Supplementary Table 11).

**Table 2.** Accuracy of 5 prediction methods in analyses of type 2 diabetes in a Latino cohort. We report adjusted *R*^2^ on the liability scale for each of the 5 main prediction methods. We obtained similar relative results using Nagelkerke *R*^2^, *R*^2^ on the observed scale and AUC (S7 Table). P-values are from likelihood ratio tests comparing models EUR and LAT to the null model, model LAT+ANC to LAT, model EUR+LAT to EUR, and EUR+LAT+ANC to EUR+LAT. For the EUR model we used *R_LD_*^2^=0.8 and *P_T_*=0.05 and for LAT we used *R_LD_*^2^=0.8 and *P_T_*=0.2. We also report normalized weights, defined as the mixing weight 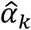 (see Methods) multiplied by the standard deviation of the PRS.

| Model | Weights associated to each predictor | | Adjusted $R^2$ | P-value for improvement over simpler model |
| --- | --- | --- | --- | --- |
|  | European PRS | Latino PRS |  |  |
| EUR | 0.16490 | | 0.02700 | $<10^{-49}$ |
| LAT | | 0.14332 | 0.02030 | $<10^{-37}$ |
| LAT+ANC | | 0.14623 | 0.03362 | $<10^{-24}$ |
| EUR+LAT | 0.16344 | 0.14164 | 0.04735 | $<10^{-37}$ |
| EUR+LAT+ANC | 0.17629 | 0.14108 | 0.04736 | 0.3 |

We investigated how the prediction accuracy of each method varied as a function of P-value thresholds, by varying either the EUR P-value threshold (Fig 2a and S12 Table) or the LAT P-value threshold (Fig 2b and S13 Table) between 10^-8^ and 1. In both cases, permissive P-value thresholds performed best, reflecting the relatively small sample sizes analyzed. However, the prediction accuracy of EUR+LAT+ANC was relatively stable, with prediction adjusted *R*^2^ > 0.037 across all EUR P-value thresholds (Fig 2a) and adjusted *R*^2^ > 0.033 across all LAT P-value thresholds (Fig 2b). In Fig 2a, we observe that as the EUR P-value threshold becomes more stringent, the difference in prediction accuracy between EUR+LAT+ANC and EUR+LAT increases, because EUR is less able to capture polygenic ancestry effects (see above).

**Fig 2.**
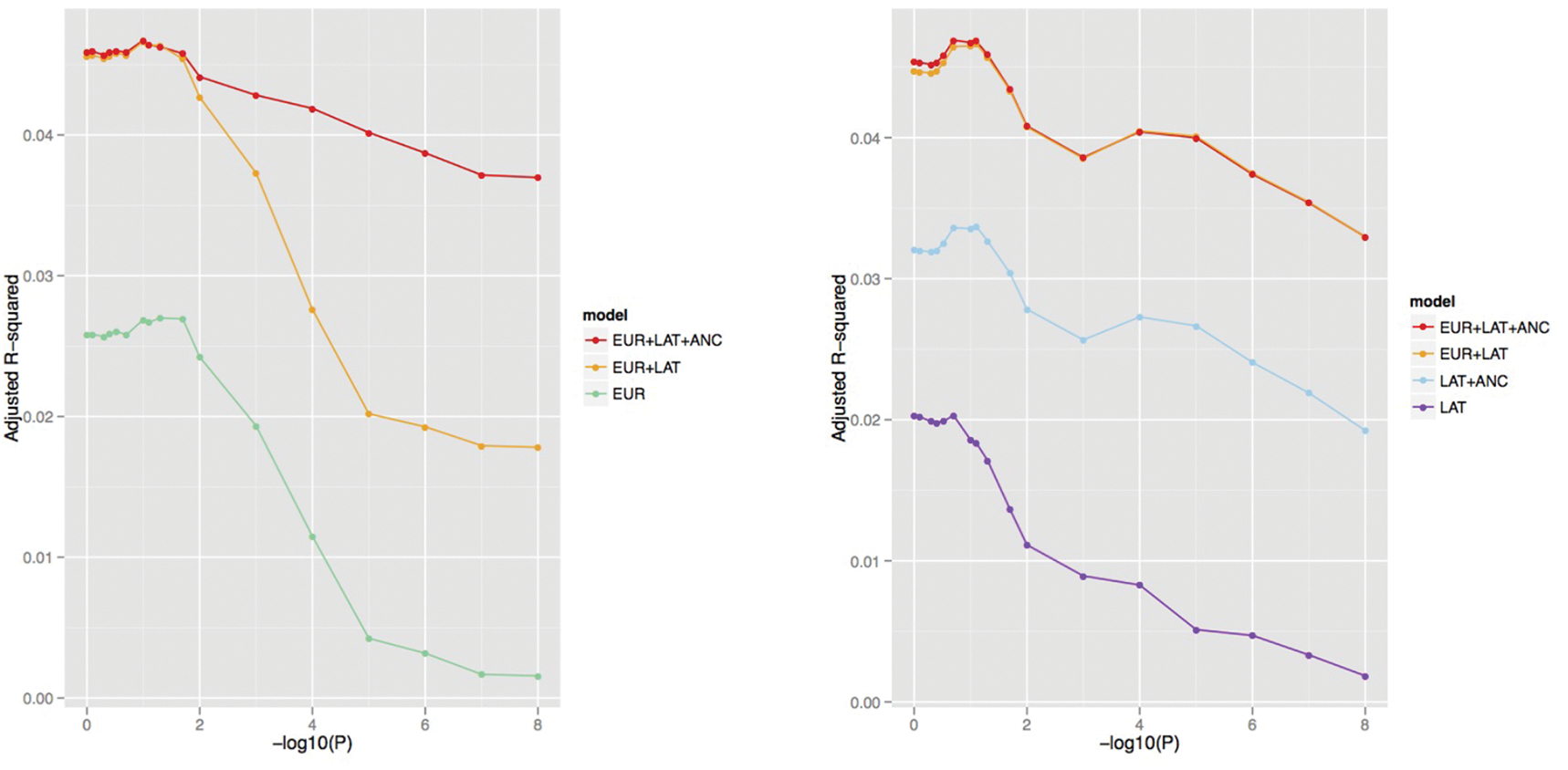
Accuracy of 5 prediction methods in analyses of type 2 diabetes in a Latino cohort as a function of P-value thresholds. We report prediction accuracies for each of the 5 main prediction methods as a function of (a) EUR P-value threshold, where applicable (with optimized LAT P-value threshold, where applicable) and (b) LAT P-value threshold, where applicable (with optimized EUR P-value threshold, where applicable). Numerical results are provided in S13 Table and S14 Table.

In the above results (Table 2 and Fig 2), we allowed each prediction method to optimize its mixing weights via an in-sample fit in the target sample. This procedure could in principle be susceptible to overfitting [34,35]. We did not expect overfitting to be a concern given the small number of mixing weights optimized (at most 3) relative to the target sample size (8,181) and given our use of adjusted *R*^2^ as the evaluation metric, but to verify this expectation, we repeated our analyses using 10x9-fold cross-validation (see Methods). Methods that use two training populations remained much more accurate than single ancestry methods, as prediction accuracy decreased only very slightly (2-4% relative decrease vs. Table 2) for each method (S14 Table). These slight decreases are expected, since mixing weights optimized within 10x9 cross-validation are slightly suboptimal (due to reduced training data) and prediction accuracy is mildly sensitive to the choice of mixing weights (S2 Fig).

### Analyses of type 2 diabetes in South Asians

We applied the same methods to predict T2D in South Asian target samples from the UK Biobank. We used publicly available European summary statistics from DIAGRAM (12,171 cases and 56,862 controls; effective sample size = 40,101) as European training data, South Asian summary statistics from SAT2D [27] (5,561 cases and 14,458 controls; effective sample size = 16,065) as South Asian training data, and UK Biobank genotypes and phenotypes (272 cases and 1,484 controls; effective sample size = 919) as South Asian validation data (see Methods).

Prediction accuracies (adjusted *R*^2^ on the liability scale [21], assuming sample prevalence 15%) and optimal weights for the 5 main methods (EUR, SAS, SAS+ANC, SAS+LAT, EUR+SAS+ANC) are reported in Table 3 (other prediction metrics are reported in S15 Table). In each case, the best prediction accuracy was obtained using LD-pruning threshold *R*_*LD*_^2^=0.8 (results using different LD-pruning thresholds are reported in S16 Table); the value of the optimal P-value threshold *P_T_* was equal to 10^−3^ for EUR and 0.8 for SAS. EUR performed only 14% better than SAS despite the larger training sample size, again reflecting a tradeoff between sample size and target-matched LD patterns. EUR+SAS attained 72%-95% relative improvements vs. EUR and SAS respectively (and used a slightly larger weight for EUR than for SAS). Adding an ancestry predictor to EUR+SAS produced an insignificant change in accuracy for EUR+ SAS +ANC compared to EUR+SAS; we note a modest correlation between each prediction method and the proportion of European-related ancestry [36] within South Asian samples (see S17 Table). We repeated our analyses using stratified 10-fold cross-validation to estimate mixing weights (see Methods). We observed that methods that use two training populations continued to substantially outperform PRS using a single training population despite a decrease in prediction adjusted *R*^2^ (vs. Table 3) for each method, consistent with the limited sample size for estimating mixing weights (S18 Table).

**Table 3.** Accuracy of 5 prediction methods in analyses of type 2 diabetes in a South Asian cohort. We report adjusted *R*^2^ on the liability scale for each of the 5 main prediction methods. We obtained similar relative results using Nagelkerke *R*^2^, *R*^2^ on the observed scale and AUC (S16 Table). P-values are from likelihood ratio tests comparing models EUR and SAS to the null model, model SAS+ANC to SAS, model EUR+SAS to EUR, and EUR+LAT+ANC to EUR+SAS. For the EUR model we used *R*_*LD*_^2^=0.8 and *P*_*T*_=10^−3^ and for SAS we used *R*_*LD*_^2^=0.8 and *P*_*T*_=0.8. We also report normalized weights, defined as the mixing weight 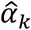 (see Methods) multiplied by the standard deviation of the PRS.

| Model | Weights associated to each predictor | | Adjusted $R^2$ | P-value for improvement over simpler model |
| --- | --- | --- | --- | --- |
|  | European PRS | SAS PRS |  |  |
| EUR | 0.09001 | | 0.01767 | $<10^{-3}$ |
| SAS | | 0.08488 | 0.01556 | $<10^{-3}$ |
| SAS+ANC |  | 0.08821 | 0.01572 | 0.28 |
| EUR+SAS | 0.08309 | 0.07746 | 0.03031 | $<10^{-2}$ |
| EUR+SAS+ANC | 0.08138 | 0.07989 | 0.02968 | 0.46 |

### Analyses of height in Africans

We applied the same methods to predict height in African target samples from the UK Biobank. We used European summary statistics from UK Biobank (113,660 samples; British ancestry only) as European training data, African summary statistics from ref. [28] (20,427 samples) as African training data, and African UK Biobank genotypes and phenotypes (1,745 samples) as African validation data.

Prediction accuracies (adjusted *R*^2^) and optimal weights for the 5 main methods (EUR, AFR, AFR+ANC, EUR+AFR, EUR+AFR+ANC) are reported in Table 4. For EUR and AFR, the best prediction accuracy was obtained using *R_LD_*^2^=0.2 and *R_LD_*^2^=0.8 respectively, thus we used these respective values of *R_LD_*^2^for EUR and AFR in each PRS in all primary analyses (results using different LD thresholds are reported in S19 Table); the value of the optimal P-value threshold *P_T_* was equal to 10^−3^ for EUR and 0.05 for AFR. EUR performed much better than AFR, consistent with the far larger training sample size.

**Table 4.** Accuracy of 5 prediction methods in analyses of height in an African cohort. We report adjusted *R*^2^ on the observed scale for each of the 5 main prediction methods. P-values are from likelihood ratio tests comparing models EUR and AFR to the null model, model AFR+ANC to AFR, model EUR+AFR to EUR, and EUR+LAT+ANC to EUR+AFR. For the EUR model we used *R_LD_*^2^=0.2 and *P_T_*=10^−3^ and for AFR we used *R_LD_*^2^=0.8 and *P_T_*=0.05. We also report normalized weights, defined as the mixing weight 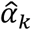 (see Methods) multiplied by the standard deviation of the PRS.

| Model | Weights associated to each predictor | | Adjusted $R^2$ | P-value for improvement over simpler model |
| --- | --- | --- | --- | --- |
|  | European PRS | African PRS |  |  |
| EUR | 0.164 | | 0.02618 | $<10^{-11}$ |
| AFR | | 0.106 | 0.01074 | $<10^{-5}$ |
| AFR+ANC |  | 0.124 | 0.01331 | 0.01 |
| EUR+AFR | 0.155 | 0.092 | 0.03397 | $<10^{-3}$ |
| EUR+AFR+ANC | 0.150 | 0.102 | 0.03443 | 0.17 |

Nevertheless, EUR+AFR attained a 30% improvement vs. EUR (using a larger weight for EUR than for AFR). Adding an ancestry predictor to EUR+AFR produced an insignificant change in accuracy for EUR+AFR+ANC compared to EUR+AFR; we note a modest correlation between each prediction method and the proportion of European-related ancestry [36] within African samples (see S20 Table). We repeated our analyses using stratified 10-fold cross-validation to estimate mixing weights (see Methods). We observed that methods that use two training populations continued to substantially outperform PRS using a single training population despite a decrease in prediction adjusted *R*^2^ (vs. Table 4) for each method, consistent with the limited sample size for estimating mixing weights (S21 Table).

## Discussion

We have shown that combining training data from European samples and training data from the target population attains a >70% relative improvement in prediction accuracy for type 2 diabetes in both Latino and South Asian cohorts compared to prediction methods that use training data from a single population. In addition, this approach attains 30% relative improvement in prediction accuracy for height in an African cohort. These relative improvements are robust to overfitting, consistent with simulations and reduce the documented gap in risk prediction accuracy between European and non-European target populations [2,8,18,19,37,38]. Intuitively, our approach leverages both large training sample sizes and training data with target-matched LD patterns. We note that the effects of differential tagging (or different causal effect sizes) in different populations can potentially be quantified using cross-population genetic correlation [39–41], and that leveraging data from a different population to improve predictions is a natural analogue to leveraging data from a correlated trait [14].

Despite these advantages, our work is subject to limitations and leaves several questions open for future exploration. First, although we have demonstrated large relative improvements in prediction accuracy, absolute prediction accuracies are currently not large enough to achieve clinical utility, which will require larger sample sizes [4,5]; our simulations suggest that multi-ethnic polygenic risk scores will continue to produce improvements at larger sample sizes (Fig 1). Second, while our focus here was on prediction without using individual-level training data, when such data is available it may be possible to attain higher prediction accuracy using methods that fit all markers simultaneously, such as Best Linear Unbiased Predictor (BLUP) methods and their extensions [10–17]. Third, our LDpred risk prediction method [8], which analyzes summary statistics in conjunction with LD information from a reference panel, is more accurate in European populations than the informed LD-pruning + P-value thresholding approach employed here; we did not employ LDpred due to the complexities of admixture-LD in analyses of admixed populations that explicitly model LD [42], but extending LDpred to handle these complexities could further improve accuracy. Fourth, we note that in our application to real phenotypes adding an ancestry predictor produced insignificant changes in prediction accuracy, primarily because ancestry effects are captured by the polygenic risk scores; adding an ancestry predictor only improves prediction when we use a stringent P-value threshold to build the polygenic risk score (Fig 2). Fifth, we have not considered here how to improve prediction accuracy in data sets with related individuals [15]. Finally, we focused our analyses on common variants, but future work may wish to consider rare variants as well.

## Acknowledgements

We are grateful to B. Vilhjalmsson and L. Liang for helpful discussions. We are grateful to G. Lettre for assistance with data from ref. [28]. This research has been conducted using the UK Biobank Resource (Application Number: 16549). This research was funded by NIH grant R01 GM105857 (A.L.P.).

## Consortia

**South Asian Type 2 Diabetes (SAT2D) Consortium.** Jaspal S Kooner, Danish Saleheen, Xueling Sim, Joban Sehmi, Weihua Zhang, Philippe Frossard, Latonya F Been, Kee-Seng Chia, Antigone S Dimas, Neelam Hassanali, Tazeen Jafar, Jeremy BM Jowett, Xinzhing Li, Venkatesan Radha, Simon D Rees, Fumihiko Takeuchi, Robin Young, Tin Aung, Abdul Basit, Manickam Chidambaram, Debashish Das, Elin Grunberg, Åsa K Hedman, Zafar I Hydrie, Muhammed Islam, Chiea-Chuen Khor, Sudhir Kowlessur, Malene M Kristensen, Samuel Liju, Wei-Yen Lim, David R Matthews, Jianjun Liu, Andrew P Morris, Alexandra C Nica, Janani M Pinidiyapathirage, Inga Prokopenko, Asif Rasheed, Maria Samuel, Nabi Shah, A Samad Shera, Kerrin S Small, Chen Suo, Ananda R Wickremasinghe, Tien Yin Wong, Mingyu Yang, Fan Zhang, DIAGRAM, MuTHER, Goncalo R Abecasis, Anthony H Barnett, Mark Caulfield, Panos Deloukas, Tim Frayling, Philippe Froguel, Norihiro Kato, Prasad Katulanda, M Ann Kelly, Junbin Liang, Viswanathan Mohan, Dharambir K Sanghera, James Scott, Mark Seielstad, Paul Z Zimmet, Paul Elliott, Yik Ying Teo, Mark I McCarthy, John Danesh, E Shyong Tai, and John C Chambers

**The SIGMA Type 2 Diabetes Consortium.** Amy L. Williams, Suzanne B. R. Jacobs, Hortensia Moreno-Macías, Alicia Huerta-Chagoya, Claire Churchouse, Carla Márquez-Luna, Humberto García-Ortíz, María José Gómez-Vázquez, Stephan Ripke, Alisa K. Manning, Benjamin Neale, David Reich, Daniel O. Stram, Juan Carlos Fernández-López, Nick Patterson, Suzanne B. R. Jacobs, Claire Churchhouse, Shuba Gopal, James A. Grammatikos, Ian C. Smith, Kevin H. Bullock, Amy A. Deik, Amanda L. Souza, Kerry A. Pierce, Clary B. Clish, Angélica Martínez-Hernández, Francisco Barajas-Olmos, Federico Centeno-Cruz, Elvia Mendoza-Caamal, Cecilia Contreras-Cubas, Cristina Revilla-Monsalve, Sergio Islas-Andrade, Emilio Córdova, Xavier Soberón, María Elena González-Villalpando, Brian E. Henderson, Kristine Monroe, Lynne Wilkens, Laurence N. Kolonel, and Loic Le Marchand, Laura Riba, María Luisa Ordóñez-Sánchez, Rosario Rodríguez-Guillén, Ivette Cruz-Bautista, Maribel Rodríguez-Torres, Linda Liliana Muñoz-Hernández, Donají Gómez, Ulises Alvirde, Olimpia Arellano, Robert C. Onofrio, Wendy M. Brodeur, Diane Gage, Jacquelyn Murphy, Jennifer Franklin, Scott Mahan, Kristin Ardlie, Andrew T. Crenshaw, Wendy Winckler, Maria L. Cortes, Noël P. Burtt, Carlos A. Aguilar-Salinas, Clicerio González-Villalpando, Jose C. Florez, Lorena Orozco, Christopher A. Haiman, Teresa Tusié-Luna, David Altshuler

## Web Resources

PLINK: https://www.cog-genomics.org/plink2.

WTCCC2 data set: http://www.wtccc.org.uk/ccc2.

SIGMA data set: http://www.type2diabetesgenetics.org.

DIAGRAM summary association statistics: http://www.diagram-consortium/org/.

UK Biobank data set: https://www.ukbiobank.ac.uk.

FastPCA (EIGENSOFT version 6.1.4): http://www.hsph.harvard.edu/alkes-price/software/.

## Supplementary Tables

**S1 Table.** Prediction accuracy of 5 prediction methods in simulations using different LD-pruning thresholds. Reported values are mean adjusted *R*^2^ and s.e. over 100 simulations.

| **Model** | **LD-pruning thresholds** |  |  |  |
| --- | --- | --- | --- | --- |
|  | **0.1** | **0.2** | **0.5** | **0.8** |
| EUR | 0.02886<br>(0.001) | 0.03089<br>(0.001) | 0.03610<br>(0.002) | 0.03927<br>(0.002) |
| LAT | 0.02268<br>(0.002) | 0.02516<br>(0.002) | 0.02845<br>(0.003) | 0.03200<br>(0.001) |
| LAT+ANC | 0.03262<br>(0.006) | 0.03486<br>(0.006) | 0.03759<br>(0.006) | 0.04115<br>(0.002) |
| EUR+LAT | 0.05020<br>(0.002) | 0.05338<br>(0.002) | 0.05984<br>(0.002) | 0.06441<br>(0.002) |
| EUR+LAT+ANC | 0.05432<br>(0.002) | 0.05739<br>(0.002) | 0.06449<br>(0.002) | 0.07053<br>(0.002) |

**S2 Table.**
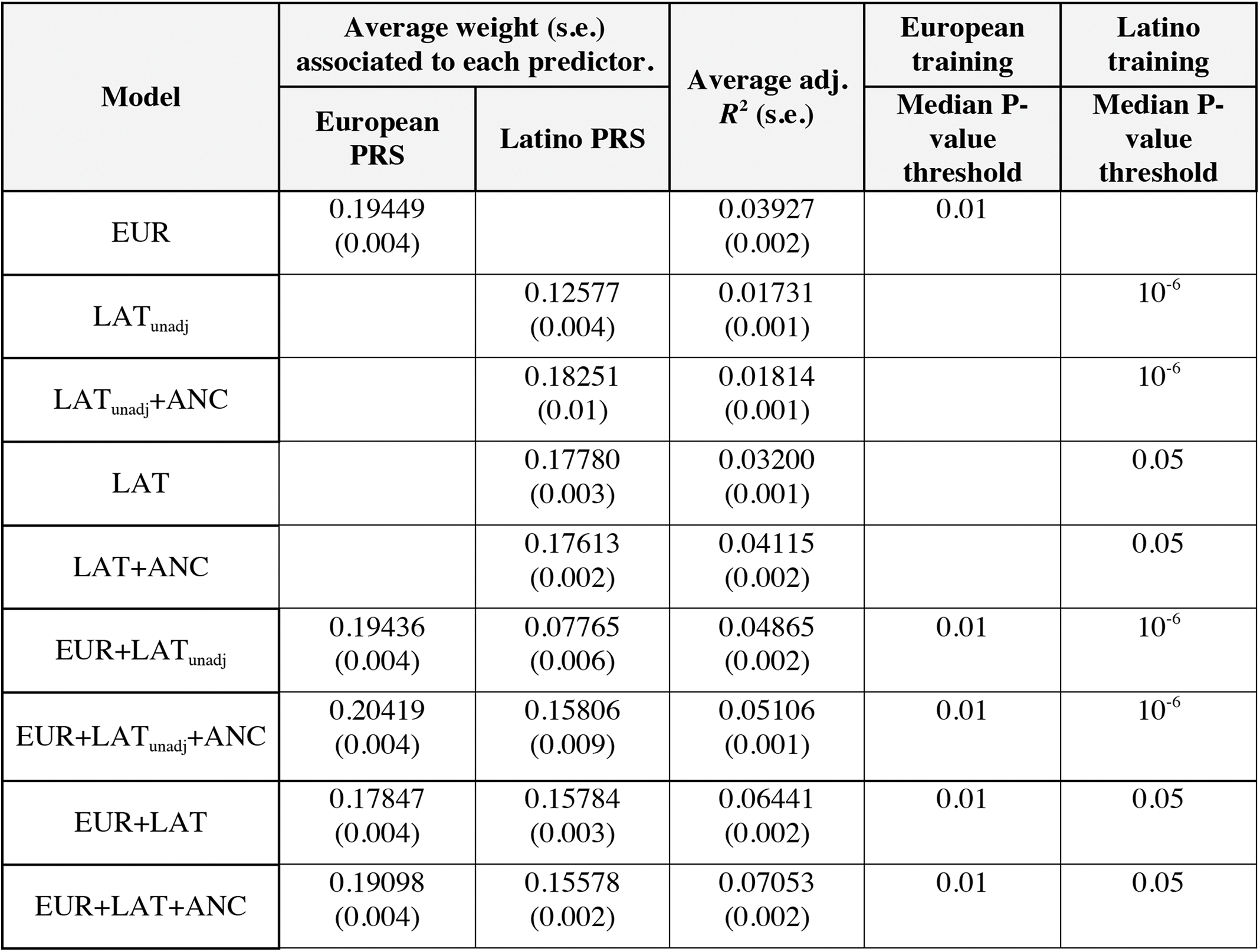
Accuracy of 9 prediction methods in simulations. We report prediction accuracies for methods using both ancestry-adjusted Latino effect sizes (LAT) and ancestry-unadjusted Latino effect sizes (LAT_unadj_). Reported values are mean atdjusted *R*^2^ over 100 simulations. We also report normalized weights, defined as the mixing weight 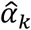 (see Methods) multiplied by the standard deviation of the PRS.

| Model | Average weight (s.e.)<br>associated to each predictor. | | Average adj.<br>$R^2$ (s.e.) | European<br>training | Latino<br>training |
| --- | --- | --- | --- | --- | --- |
|  | European<br>PRS | Latino PRS |  | Median P-value<br>threshold | Median P-value<br>threshold |
| EUR | 0.19449<br>(0.004) |  | 0.03927<br>(0.002) | 0.01 |  |
| LAT<sub>unadj</sub> | | 0.12577<br>(0.004) | 0.01731<br>(0.001) | | $10^{-6}$ |
| LAT<sub>unadj</sub>+ANC | | 0.18251<br>(0.01) | 0.01814<br>(0.001) | | $10^{-6}$ |
| LAT |  | 0.17780<br>(0.003) | 0.03200<br>(0.001) |  | 0.05 |
| LAT+ANC |  | 0.17613<br>(0.002) | 0.04115<br>(0.002) |  | 0.05 |
| EUR+LAT<sub>unadj</sub> | 0.19436<br>(0.004) | 0.07765<br>(0.006) | 0.04865<br>(0.002) | 0.01 | $10^{-6}$ |
| EUR+LAT<sub>unadj</sub>+ANC | 0.20419<br>(0.004) | 0.15806<br>(0.009) | 0.05106<br>(0.001) | 0.01 | $10^{-6}$ |
| EUR+LAT | 0.17847<br>(0.004) | 0.15784<br>(0.003) | 0.06441<br>(0.002) | 0.01 | 0.05 |
| EUR+LAT+ANC | 0.19098<br>(0.004) | 0.15578<br>(0.002) | 0.07053<br>(0.002) | 0.01 | 0.05 |

**S3 Table.** *R*^2^ with European ancestry for 9 prediction methods in simulations. European ancestry is represented by PC1 in the SIGMA data set. Reported values are mean *R*^2^ over 100 simulations. The average *R*^2^ between ancestry and phenotype was 0.011.

| Model | Average $R^2$ (s.e.) |
| --- | --- |
| EUR | 0.0254<br>(0.019) |
| LAT<sub>unadj</sub> | 0.3721<br>(0.034) |
| LAT<sub>unadj</sub>+ANC | 0.2205<br>(0.037) |
| LAT | 0.0015<br>(0.007) |
| LAT+ANC | 0.0437<br>(0.025) |
| EUR+LAT<sub>unadj</sub> | 0.0626<br>(0.02) |
| EUR+LAT<sub>unadj</sub>+ANC | 0.0337<br>(0.019) |
| EUR+LAT | 0.0103<br>(0.016) |
| EUR+LAT+ANC | 0.0178<br>(0.018) |

**S4 Table.** Accuracy of 9 prediction methods in simulations with ancestry-correlated phenotypes. We report prediction accuracies for methods using both ancestry-adjusted Latino effect sizes (LAT) and ancestry-unadjusted Latino effect sizes (LAT_unadj_). Reported values are mean adjusted *R*^2^ and s.e. over 100 simulations. We also report normalized weights, defined as the mixing weight 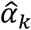 (see Methods) multiplied by the standard deviation of the PRS.

| Model | Average weight (s.e.) associated to each predictor. | | Average adj. $R^2$ (s.e.) | European training | Latino training |
| --- | --- | --- | --- | --- | --- |
|  | European PRS | Latino PRS |  | Median P-value threshold | Median P-value threshold |
| EUR | 0.19452<br>(0.004) |  | 0.03927<br>(0.002) | 0.01 |  |
| LAT<sub>unadj</sub> | | 0.01353<br>(0.011) | 0.01181<br>(0.001) | | $10^{-6}$ |
| LAT<sub>unadj</sub>+ANC | | 0.24467<br>(0.016) | 0.01359<br>(0.001) | | $10^{-6}$ |
| LAT |  | 0.17866<br>(0.002) | 0.03227<br>(0.001) |  | 0.05 |
| LAT+ANC |  | 0.17650<br>(0.002) | 0.04095<br>(0.002) |  | 0.05 |
| EUR+LAT<sub>unadj</sub> | 0.20402<br>(0.004) | 0.01035<br>(0.009) | 0.04587<br>(0.002) | 0.01 | $10^{-6}$ |
| EUR+LAT<sub>unadj</sub>+ANC | 0.20671<br>(0.004) | 0.19082<br>(0.014) | 0.04760<br>(0.002) | 0.01 | $10^{-6}$ |
| EUR+LAT | 0.17729<br>(0.004) | 0.15818<br>(0.002) | 0.06426<br>(0.002) | 0.01 | 0.05 |
| EUR+LAT+ANC | 0.19060<br>(0.004) | 0.15681<br>(0.002) | 0.06960<br>(0.002) | 0.01 | 0.05 |

**S5 Table.** Numerical values of results displayed in Fig 1. We report prediction accuracies for each of the 5 main prediction methods, for each subset of chromosomes. Reported values are mean adjusted *R*^2^ and s.e over 100 simulations.

| Model | Chr 1 | Chr 1-2 | Chr 1-4 | Chr 1-22 |
| --- | --- | --- | --- | --- |
| EUR | 0.18641 (0.003) | 0.15778 (0.003) | 0.12453 (0.002) | 0.03927 (0.002) |
| LAT | 0.14580 (0.003) | 0.11512 (0.003) | 0.08360 (0.002) | 0.03200 (0.001) |
| LAT+ANC | 0.14941 (0.003) | 0.11859 (0.003) | 0.08651 (0.002) | 0.04115 (0.002) |
| EUR+LAT | 0.21298 (0.003) | 0.18374 (0.003) | 0.14931 (0.002) | 0.06441 (0.002) |
| EUR+LAT+ANC | 0.21576 (0.003) | 0.18695 (0.003) | 0.15244 (0.002) | 0.07053 (0.002) |

**S6 Table.** Numerical values of results displayed in S1 Fig. We report prediction accuracies for each of the 5 main prediction methods, for each subset of chromosomes, in simulations including the causal SNPs.

| Model | Chr 1 | Chr 1-2 | Chr 1-4 | Chr 1-22 |
| --- | --- | --- | --- | --- |
| EUR | 0.277 (0.003) | 0.247 (0.003) | 0.207 (0.002) | 0.079 (0.003) |
| LAT | 0.143 (0.003) | 0.130 (0.003) | 0.113 (0.002) | 0.042 (0.001) |
| LAT+ANC | 0.158 (0.003) | 0.141 (0.003) | 0.120 (0.002) | 0.052 (0.002) |
| EUR+LAT | 0.295 (0.003) | 0.267 (0.003) | 0.232 (0.002) | 0.106 (0.002) |
| EUR+LAT+ANC | 0.301 (0.002) | 0.275 (0.003) | 0.243 (0.002) | 0.122 (0.002) |

**S7 Table.** Accuracy of 5 prediction methods in analyses of type 2 diabetes in a Latino cohort, using alternate prediction metrics. Liability-scale adjusted *R*^2^ was computed assuming a disease prevalence of *K*=0.08.

| Model | Observed-scale adj. $R^2$ | Liability-scale adj. $R^2$ | Nagelkerke $R^2$ | AUC |
| --- | --- | --- | --- | --- |
| EUR | 0.02707 | 0.02700 | 0.03633 | 0.59012 |
| LAT | 0.02042 | 0.02030 | 0.02742 | 0.58175 |
| LAT+ANC | 0.03361 | 0.03362 | 0.04517 | 0.60342 |
| EUR+LAT | 0.04702 | 0.04735 | 0.06311 | 0.62375 |
| EUR+LAT+ANC | 0.04703 | 0.04736 | 0.06328 | 0.62416 |

**S8 Table.**
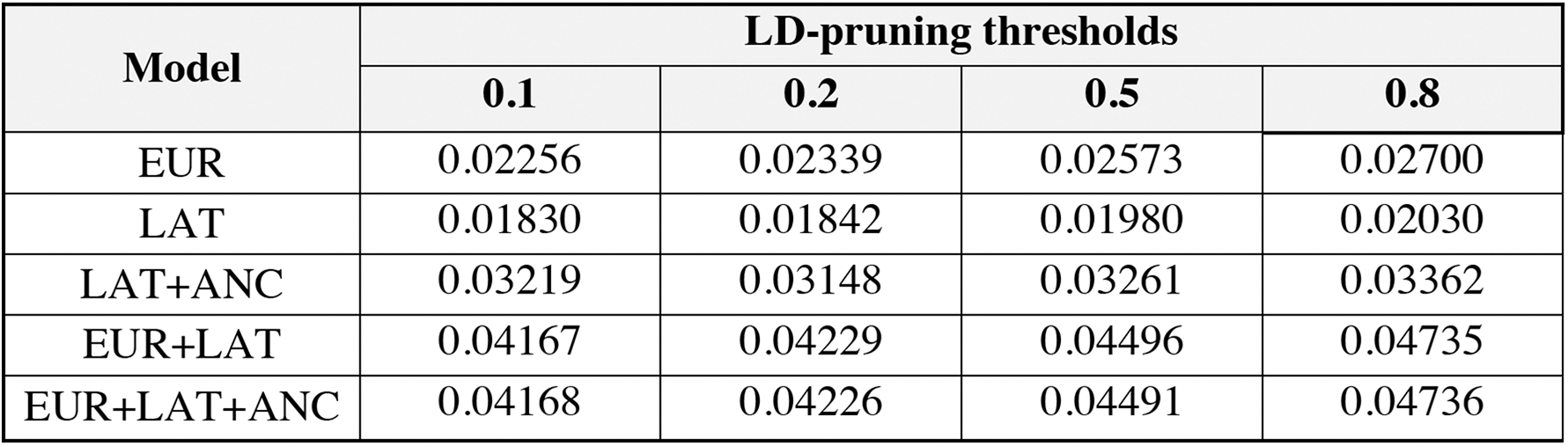
Prediction accuracy of 5 prediction methods in analyses of type 2 diabetes in a Latino cohort using different LD-pruning thresholds. Liability-scale adjusted *R*^2^ was computed assuming a disease prevalence of *K*=0.08.

| Model | LD-pruning thresholds |  |  |  |
| --- | --- | --- | --- | --- |
|  | 0.1 | 0.2 | 0.5 | 0.8 |
| EUR | 0.02256 | 0.02339 | 0.02573 | 0.02700 |
| LAT | 0.01830 | 0.01842 | 0.01980 | 0.02030 |
| LAT+ANC | 0.03219 | 0.03148 | 0.03261 | 0.03362 |
| EUR+LAT | 0.04167 | 0.04229 | 0.04496 | 0.04735 |
| EUR+LAT+ANC | 0.04168 | 0.04226 | 0.04491 | 0.04736 |

**S9 Table.** *R* and *R*^2^ with European ancestry for 9 prediction methods and T2D phenotype in analyses of type 2 diabetes in a Latino cohort. European ancestry is represented by PC1 in the SIGMA data set.

| **Model** | **$R$** | **$R^2$** |
| --- | --- | --- |
| EUR | -0.751 | 0.564 |
| LAT<sub>unadj</sub> | -0.995 | 0.990 |
| LAT<sub>unadj</sub>+ANC | -0.999 | 0.999 |
| LAT | 0.025 | 0.001 |
| LAT+ANC | -0.607 | 0.369 |
| EUR+LAT<sub>unadj</sub> | -0.684 | 0.468 |
| EUR+LAT<sub>unadj</sub><br>+ANC | -0.671 | 0.450 |
| EUR+LAT | -0.548 | 0.300 |
| EUR+LAT+ANC | -0.513 | 0.263 |
| T2D phenotype | -0.112 | 0.013 |

**S10 Table.**
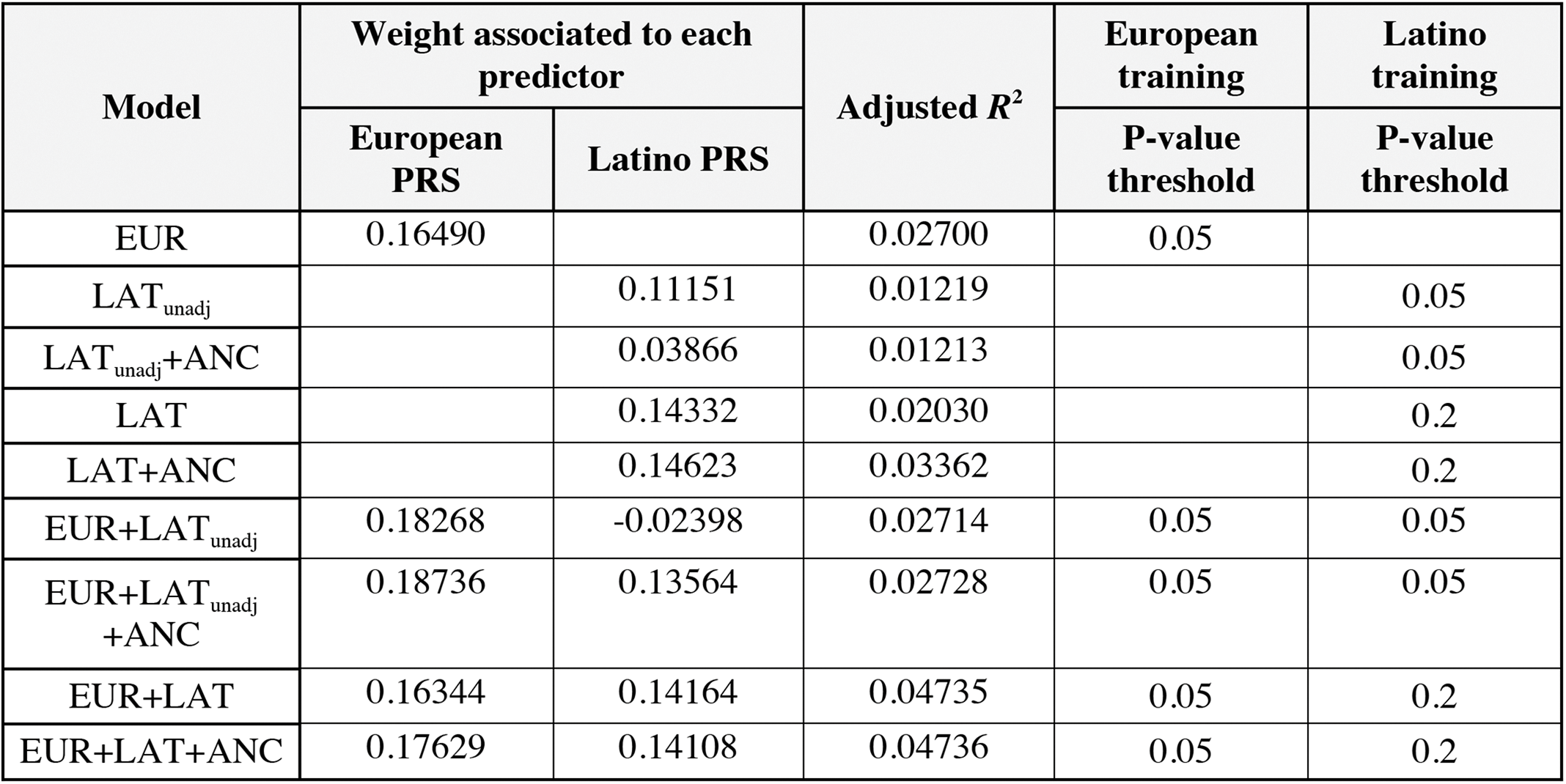
Accuracy of 9 prediction methods in analyses of type 2 diabetes in a Latino cohort. We report adjusted *R*^2^ on the liability scale for methods using both ancestry-adjusted Latino effect sizes (LAT) and ancestry-unadjusted Latino effect sizes (LAT_unadj_). We also report normalized weights, defined as the mixing weight 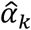 (see Methods) multiplied by the standard deviation of the PRS. We also report normalized weights, defined as the mixing weight 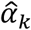 (see Methods) multiplied by the standard deviation of the PRS.

| Model | Weight associated to each predictor | | Adjusted $R^2$ | European training | Latino training |
| --- | --- | --- | --- | --- | --- |
|  | European PRS | Latino PRS |  | P-value threshold | P-value threshold |
| EUR | 0.16490 |  | 0.02700 | 0.05 |  |
| LAT<sub>unadj</sub> |  | 0.11151 | 0.01219 |  | 0.05 |
| LAT<sub>unadj</sub>+ANC |  | 0.03866 | 0.01213 |  | 0.05 |
| LAT |  | 0.14332 | 0.02030 |  | 0.2 |
| LAT+ANC |  | 0.14623 | 0.03362 |  | 0.2 |
| EUR+LAT<sub>unadj</sub> | 0.18268 | -0.02398 | 0.02714 | 0.05 | 0.05 |
| EUR+LAT<sub>unadj</sub>+ANC | 0.18736 | 0.13564 | 0.02728 | 0.05 | 0.05 |
| EUR+LAT | 0.16344 | 0.14164 | 0.04735 | 0.05 | 0.2 |
| EUR+LAT+ANC | 0.17629 | 0.14108 | 0.04736 | 0.05 | 0.2 |

**S11 Table.**
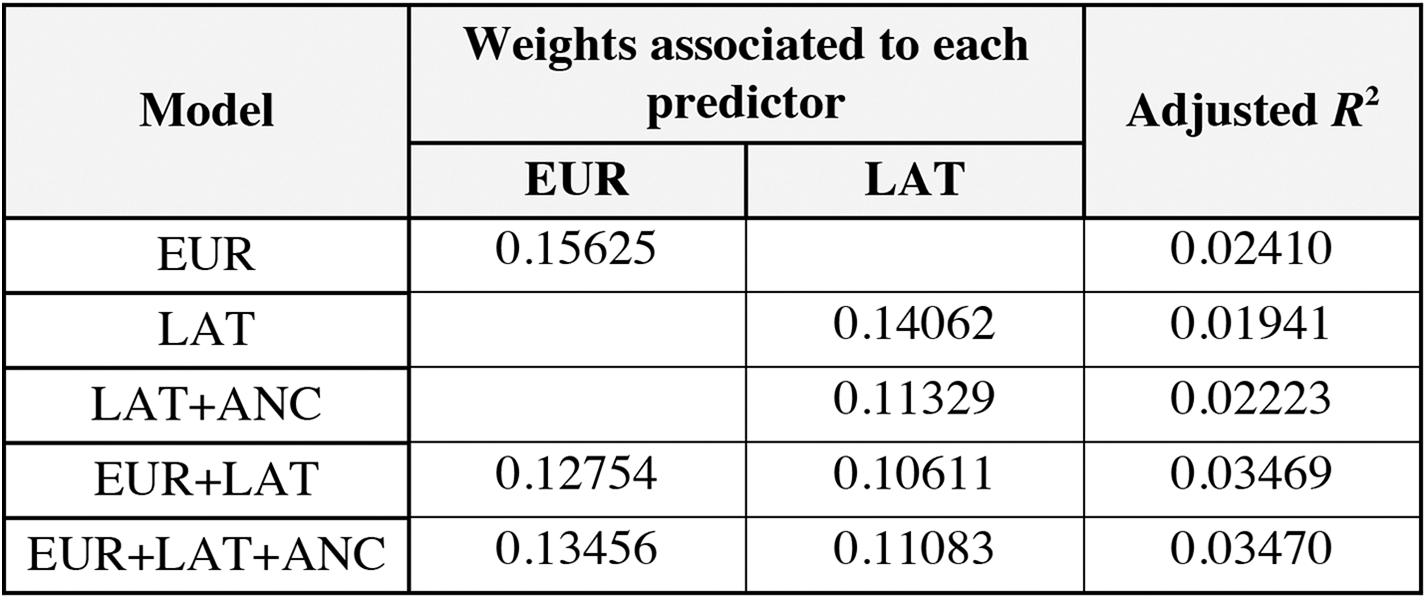
Accuracy of 5 prediction methods in analyses of type 2 diabetes in a Latino cohort using imputed genotypes. We report *R*^2^ on the liability scale for each of the 5 main prediction methods. We also report normalized weights, defined as the mixing weight 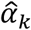 (see Methods) multiplied by the standard deviation of the PRS.

| Model | Weights associated to each predictor | | Adjusted $R^2$ |
| --- | --- | --- | --- |
|  | EUR | LAT |  |
| EUR | 0.15625 |  | 0.02410 |
| LAT |  | 0.14062 | 0.01941 |
| LAT+ANC |  | 0.11329 | 0.02223 |
| EUR+LAT | 0.12754 | 0.10611 | 0.03469 |
| EUR+LAT+ANC | 0.13456 | 0.11083 | 0.03470 |

**S12 Table.**
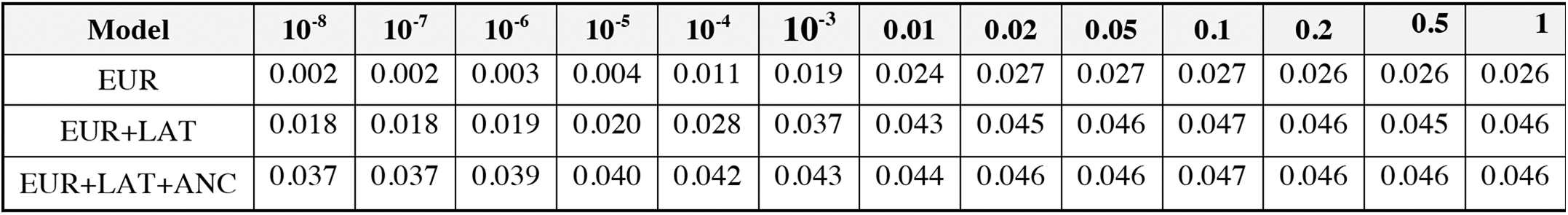
Numerical values for results displayed in Fig 2a. We report prediction adjusted *R*^2^ for each of the 3 prediction methods that include the EUR predictor.

| Model | $10^{-8}$ | $10^{-7}$ | $10^{-6}$ | $10^{-5}$ | $10^{-4}$ | $10^{-3}$ | 0.01 | 0.02 | 0.05 | 0.1 | 0.2 | 0.5 | 1 |
| --- | --- | --- | --- | --- | --- | --- | --- | --- | --- | --- | --- | --- | --- |
| EUR | 0.002 | 0.002 | 0.003 | 0.004 | 0.011 | 0.019 | 0.024 | 0.027 | 0.027 | 0.027 | 0.026 | 0.026 | 0.026 |
| EUR+LAT | 0.018 | 0.018 | 0.019 | 0.020 | 0.028 | 0.037 | 0.043 | 0.045 | 0.046 | 0.047 | 0.046 | 0.045 | 0.046 |
| EUR+LAT+ANC | 0.037 | 0.037 | 0.039 | 0.040 | 0.042 | 0.043 | 0.044 | 0.046 | 0.046 | 0.047 | 0.046 | 0.046 | 0.046 |

**S13 Table.** Numerical values for results displayed in Fig 2b. We report prediction adjusted *R*^2^ for each of the 4 prediction methods that include the LAT predictor.

|  | P-value Threshold |  |  |  |  |  |  |  |  |  |  |  |  |
| --- | --- | --- | --- | --- | --- | --- | --- | --- | --- | --- | --- | --- | --- |
| Model | $10^{-8}$ | $10^{-7}$ | $10^{-6}$ | $10^{-5}$ | $10^{-4}$ | $10^{-3}$ | 0.01 | 0.02 | 0.05 | 0.1 | 0.2 | 0.5 | 1 |
| LAT | 0.002 | 0.003 | 0.005 | 0.005 | 0.008 | 0.009 | 0.011 | 0.014 | 0.017 | 0.019 | 0.020 | 0.020 | 0.020 |
| LAT+ANC | 0.019 | 0.022 | 0.024 | 0.027 | 0.027 | 0.026 | 0.028 | 0.030 | 0.033 | 0.034 | 0.034 | 0.032 | 0.032 |
| EUR+LAT | 0.033 | 0.035 | 0.037 | 0.040 | 0.040 | 0.039 | 0.041 | 0.043 | 0.046 | 0.046 | 0.046 | 0.045 | 0.045 |
| EUR+LAT+ANC | 0.033 | 0.035 | 0.037 | 0.040 | 0.040 | 0.039 | 0.041 | 0.043 | 0.046 | 0.047 | 0.047 | 0.045 | 0.045 |

**S14 Table.** Accuracy of 5 prediction methods in analyses of type 2 diabetes in a Latino cohort, using 10x9-fold cross-validation. We report adjusted *R*^2^ on the liability scale for each of the 5 main prediction methods, and the average of adjusted *R*^2^ within each fold. Adjusted *R*^2^ merging folds is lower than average adjusted *R*^2^ across folds because of miscalibration between folds. We used 10-fold cross-validation for EUR and 10x9-fold cross-validation for LAT, LAT+ANC, EUR+LAT and EUR+LAT+ANC (see Methods). We also report normalized weights, defined as the mixing weight 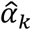 (see Methods) multiplied by the standard deviation of the PRS.

| Model | Average weight (s.d.) associated to each predictor. | | Avg. adj. $R^2$ across folds (s.d.) | Adj. $R^2$ merging folds |
| --- | --- | --- | --- | --- |
|  | European PRS | Latino PRS |  |  |
| EUR | 0.165<br>(0.004) |  | 0.02731<br>(0.014) | 0.02650 |
| LAT |  | 0.133<br>(0.012) | 0.01966<br>(0.006) | 0.01997 |
| LAT+ANC |  | 0.130<br>(0.008) | 0.03230<br>(0.009) | 0.03267 |
| EUR+LAT | 0.158<br>(0.008) | 0.125<br>(0.008) | 0.04645<br>(0.014) | 0.04646 |
| EUR+LAT+ANC | 0.177<br>(0.008) | 0.125<br>(0.008) | 0.04596<br>(0.014) | 0.04593 |

**S15 Table.** Accuracy of 5 prediction methods in analyses of type 2 diabetes in a South Asian cohort, using alternate prediction metrics. Liability-scale adjusted *R*^2^ was computed using the sample disease prevalence estimate of *K*=0.15.

| Model | Observed-scale adj. $R^2$ | Liability-scale adj. $R^2$ | Nagelkerke $R^2$ | AUC |
| --- | --- | --- | --- | --- |
| EUR | 0.00753 | 0.01767 | 0.01423 | 0.57453 |
| SAS | 0.00664 | 0.01556 | 0.01243 | 0.55606 |
| SAS+ANC | 0.00670 | 0.01572 | 0.01359 | 0.56153 |
| EUR+SAS | 0.01292 | 0.03031 | 0.02454 | 0.59155 |
| EUR+SAS+ANC | 0.01265 | 0.02968 | 0.02507 | 0.59366 |

**S16 Table.** Prediction accuracy of 5 prediction methods in analyses of type 2 diabetes in a South Asian cohort using different LD-pruning thresholds. Liability-scale adjusted *R*^2^ was computed using the sample disease prevalence estimate of *K*=0.15.

| Model | LD-pruning threshold |  |  |  |
| --- | --- | --- | --- | --- |
|  | 0.1 | 0.2 | 0.5 | 0.8 |
| EUR | 0.01064 | 0.01272 | 0.01380 | 0.01767 |
| SAS | 0.01212 | 0.00994 | 0.01196 | 0.01556 |
| SAS+ANC | 0.01220 | 0.01000 | 0.01203 | 0.01572 |
| EUR+SAS | 0.02213 | 0.02209 | 0.02456 | 0.03031 |
| EUR+SAS+ANC | 0.02157 | 0.02120 | 0.02366 | 0.02968 |

**S17 Table.** *R* and *R*^2^ with European ancestry for 5 prediction methods and T2D phenotype in analyses of type 2 diabetes in a South Asian cohort. European ancestry is represented by PC1 in the data set.

| Model | $R$ with PC1 | $R^2$ with PC1 |
| --- | --- | --- |
| EUR | -0.08572 | 0.00735 |
| SAS | 0.13099 | 0.01716 |
| SAS+ANC | -0.15702 | 0.02466 |
| EUR+SAS | 0.02550 | 0.00065 |
| EUR+SAS+ANC | -0.11607 | 0.01347 |
| T2D phenotype | -0.01390 | 0.00019 |

**S18 Table.** Accuracy of 5 prediction methods in analyses of type 2 diabetes in a South Asian cohort, using stratified 10-fold cross-validation. We report adjusted *R*^2^ on the liability scale averaged over 500 different partitions of the data into 10 stratified folds, and the average of adjusted *R*^2^ within each fold. Adjusted *R*^2^ merging folds is lower than average adjusted *R*^2^ across folds because of miscalibration between folds. We used 10-fold cross-validation for all methods, including EUR and SAS (see Methods). We also report normalized weights, defined as the mixing weight 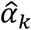 (see Methods) multiplied by the standard deviation of the PRS.

| Model | Weight<br>EUR PRS | Weight<br>SAS PRS | Avg. adj.<br>$R^2$ across<br>folds<br>(s.d) | Adj. $R^2$<br>merging<br>folds |
| --- | --- | --- | --- | --- |
| EUR | 0.09001<br>(0.007) |  | 0.01681<br>(0.031) | 0.01519 |
| SAS |  | 0.08487<br>(0.008) | 0.01700<br>(0.035) | 0.01257 |
| SAS+ANC |  | 0.08821<br>(0.008) | 0.01572<br>(0.034) | 0.01188 |
| EUR+SAS | 0.08310<br>(0.007) | 0.07745<br>(0.008) | 0.02785<br>(0.039) | 0.02614 |
| EUR+SAS+ANC | 0.08140<br>(0.007) | 0.07987<br>(0.008) | 0.02642<br>(0.039) | 0.02462 |

**S19 Table.** Prediction accuracy of 5 prediction methods in analyses of height in an African cohort using different LD-pruning thresholds.

| Model | LD-pruning threshold |  |  |  |
| --- | --- | --- | --- | --- |
|  | 0.1 | 0.2 | 0.5 | 0.8 |
| EUR | 0.01442 | 0.02619 | 0.02215 | 0.02235 |
| AFR | 0.00785 | 0.00877 | 0.01023 | 0.01075 |
| AFR+ANC | 0.00981 | 0.01081 | 0.01238 | 0.01332 |
| EUR+AFR | 0.02095 | 0.03319 | 0.03103 | 0.02940 |
| EUR+AFR+ANC | 0.02420 | 0.03344 | 0.03048 | 0.03019 |

**S20 Table.** *R* and *R*^2^ with European ancestry for 5 prediction methods and height phenotype in analyses of height in an African cohort. European ancestry is represented by PC1 in the data set.

| **Model** | **$R$ with PC1** | **$R^2$ with PC1** |
| --- | --- | --- |
| EUR | -0.12249 | 0.01500 |
| AFR | 0.29584 | 0.08752 |
| AFR+ANC | -0.18300 | 0.03349 |
| EUR+AFR | 0.04358 | 0.00190 |
| EUR+AFR+ANC | -0.11575 | 0.01340 |
| Height | -0.02199 | 0.00048 |

**S21 Table.**
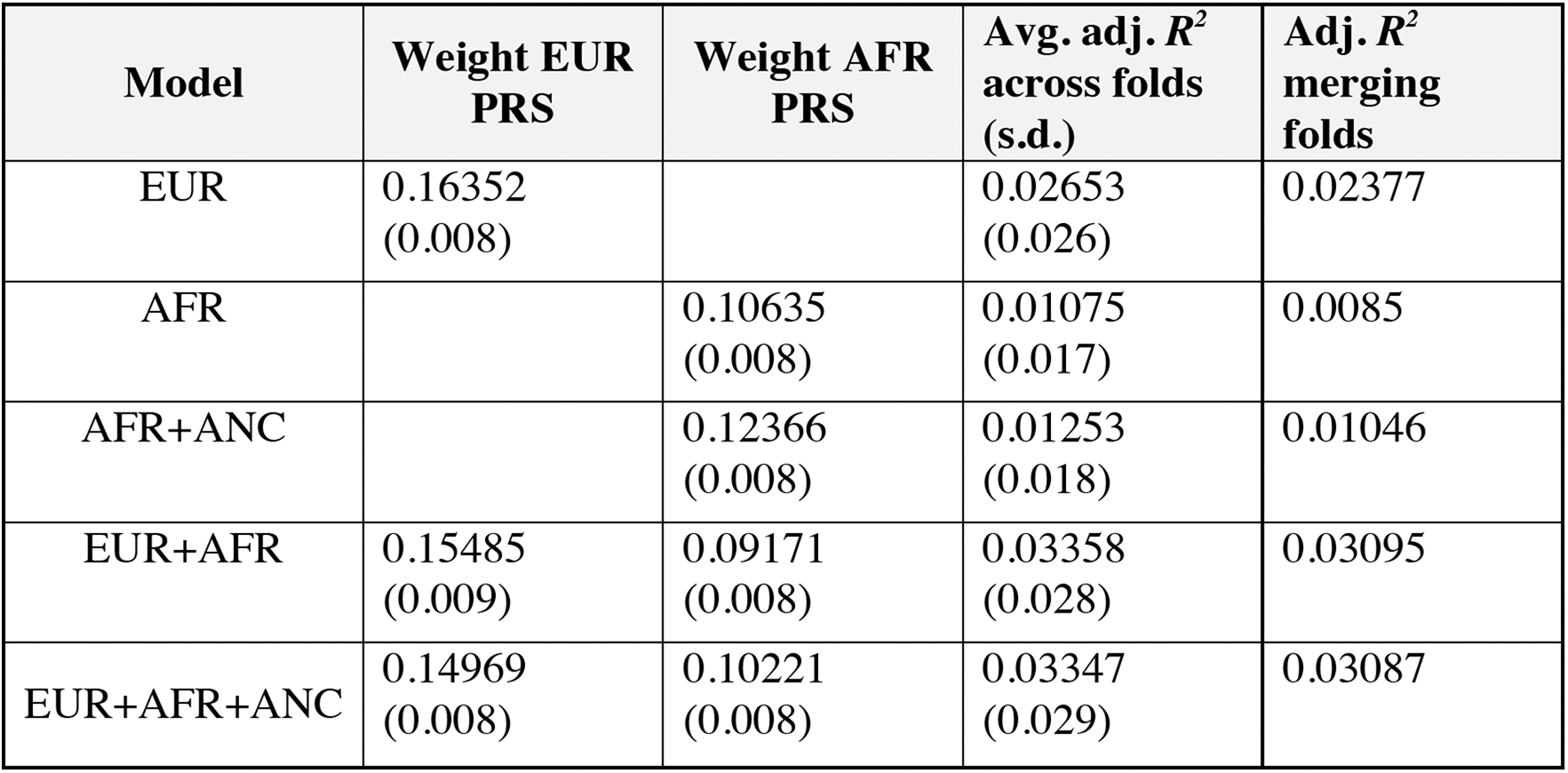
Accuracy of 5 prediction methods in analyses of height in an African cohort, using 10-fold cross validation. We report adjusted *R*^2^ merging folds averaged over 500 different partitions of the data into 10 stratified folds, and the average of adjusted *R*^2^ within each fold. Adjusted *R*^2^ merging folds is lower than average adjusted *R*^2^ across folds because of miscalibration between folds. We used 10-fold cross-validation for all methods, including EUR and AFR (see Methods). We also report normalized weights, defined as the mixing weight 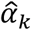 (see Methods) multiplied by the standard deviation of the PRS.

| Model | Weight EUR<br>PRS | Weight AFR<br>PRS | Avg. adj. $R^2$<br>across folds<br>(s.d.) | Adj. $R^2$<br>merging<br>folds |
| --- | --- | --- | --- | --- |
| EUR | 0.16352<br>(0.008) |  | 0.02653<br>(0.026) | 0.02377 |
| AFR |  | 0.10635<br>(0.008) | 0.01075<br>(0.017) | 0.0085 |
| AFR+ANC |  | 0.12366<br>(0.008) | 0.01253<br>(0.018) | 0.01046 |
| EUR+AFR | 0.15485<br>(0.009) | 0.09171<br>(0.008) | 0.03358<br>(0.028) | 0.03095 |
| EUR+AFR+ANC | 0.14969<br>(0.008) | 0.10221<br>(0.008) | 0.03347<br>(0.029) | 0.03087 |

## Supplementary Figures

**S1 Fig.**
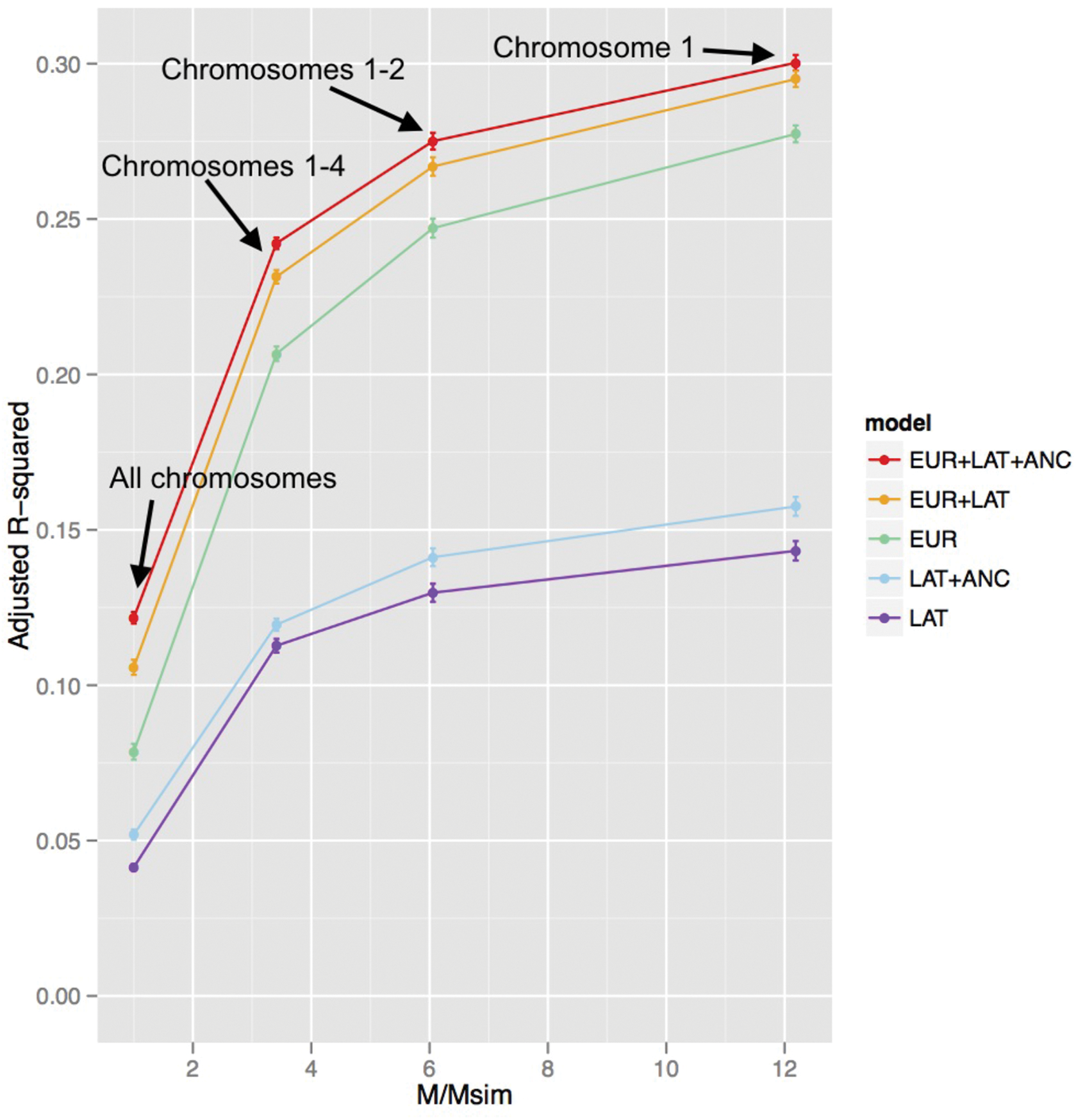
Accuracy of 5 prediction methods in simulations using subsets of chromosomes, including the causal SNPs. We report prediction accuracies for each of the 5 main prediction methods as a function of M/Msim, where M=232,629 is the total number of SNPs and Msim is the actual number of SNPS used in each simulation: 232,629 (all chromosomes), 68,188 (chromosomes 1-4), 38,412 (chromosomes 1-2), and 19,087 (chromosome 1). Numerical results are provided in S6 Table.

**S2 Fig.**
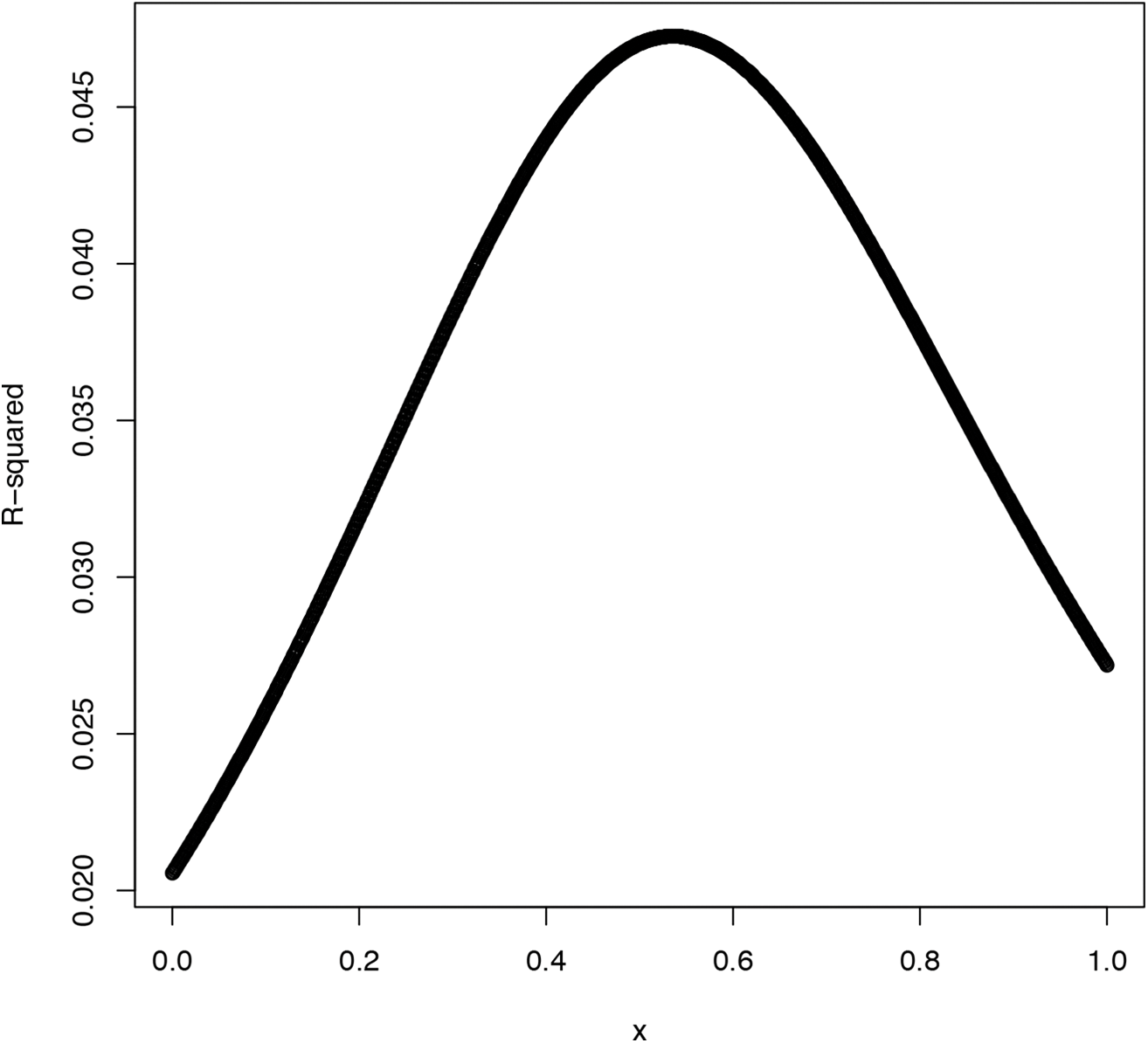
Sensitivity to mixing weights in analyses of type 2 diabetes in a Latino cohort. We report the prediction *R*^2^ of xEUR + (1-x)LAT, with x varying between 0 and 1.

